# Characterization of the cellular and subcellular distribution of fms-like tyrosine kinase 3 (FLT3) and other neuronal proteins using an alkaline phosphatase (AP) immunolabeling method

**DOI:** 10.1101/2024.10.02.616387

**Authors:** Yuqin Yin, Kathleen He, Jane Kirby, Ishraq Harque, Xin Tang

## Abstract

Precisely localizing the spatial distribution of proteins within various cell types in the brain, and in sub-cellular compartments such as the synapses, is critical for generating and testing hypothesis to elucidating the function of target proteins in the brain. The fms-like tyrosine kinase-3 (FLT3) has been studied extensively in the context of blood cell development and leukemia pathogenesis, but little is known about its role in the brain. Previous work characterizing FLT3 protein expression in the brain have not yielded convincing results, mainly limited by the low expression level of FLT3 and poor sensitivity of standard immunolabeling method using fluorescent secondary antibody. In this study, we report the systematic characterization of FLT3 protein localization during brain development using a highly sensitive immunolabeling methodology based on alkaline phosphatase (AP) polymer biochemistry. Our results reveal a previously unknown neuron-selective FLT3 expression pattern in both mouse and human cerebellum tissue samples, and demonstrate a gradual increase in the total FLT3 protein level and a cytosolic-to-dendritic change in subcellular FLT3 distribution during mouse cerebellum development. Through combining the AP immunolabeling of FLT3 with standard immunostaining of various cell type markers to achieve hybrid co-labeling of multiple antigens in tissue sections, we demonstrate that the main cell type that express FLT3 in the cerebellum is PV+, Calbindin+ Purkinje cells. To expand the use case of AP immunolabeling method in labeling neuronal proteins, we show robust labeling of Kir2.1, a potassium channel protein that is expressed at low level in neurons, in brain tissue sections collected from mouse, pig, and human brains. We further established that the AP immunolabeling method can be used in human stem cell-derived neurons to detect postsynaptic density scaffold protein PSD95 within fine subcellular structures such as dendritic spines and synapses, in cultured primary mouse cortical neurons and human stem cell-derived neurons. To our knowledge, our work represents the first report of applying AP immunolabeling to detect neuronal protein expression in brain tissue and cell samples. Moreover, our work reveals a previously unknown neuron-specific pattern of FLT3 expression in the brain, providing the foundation for further mechanistic studies to uncover its role in brain development and functioning.

## Introduction

The brain expresses the majority of the known protein-coding genes^*1*^. However, for a large number brain-expressed genes including signaling receptors, ion channels, transporters, and synaptic proteins, the cellular expression and subcellular distribution patterns of their protein products have not been well established. One such example is the receptor tyrosine kinase FLT3, which plays a critical role in the development of blood cells and its dysregulation leads to the pathogenesis of leukemia^2^. The presence of FLT3 mRNA and protein in the brain has been reported in previous studies but have not been rigorously examined^3, 4^. Results from single cell sequencing of mouse and human brain tissue indicate a high level of FLT3 mRNA expression in the cerebellum tissue, especially in the Purkinje cells^*5*^. However, it remains unknown the extent to which the expression pattern of FLT3 protein correlates with its mRNA level in cells.

Moreover, the subcellular localization of the kinase protein plays an important role in determining signaling mode and strength^*6*^. Such information can not be inferred from RNA sequencing. Our group’s previous unbiased drug screening work discovers that pharmacological inhibition of FLT3 kinase signaling in brain cells enhances the expression of KCC2, a chloride transporter protein that is required for GABAergic inhibition and is exclusively expressed in neurons^*7*^. These results suggest a previously unknown role of FLT3 signaling in neurons, and highlights the need for an *in situ* protein detection method to study FLT3 in the brain tissue.

We therefore seek to develop a immunolabeling method with high sensitivity and resolution to investigate FLT3 expression pattern in neurons. Standard fluorescent immunostaining method mainly relies on fluorescent secondary antibodies which are large molecular size proteins that has limited binding affinity, signal amplification, and sensitivity. An alternative immunolabeling often employed in histology analysis involves attaching the catalytic HRP (Horseradish peroxidase) enzyme to secondary antibodies to facilitate an *in situ* reaction to oxidize a chromogenic substrate, resulting in robust signal amplification. However, the HRP method produces a monochromatic deposit that lacks fluorescence and interferes with fluorescent light emission, thereby precluding the simultaneous staining of target proteins with other fluorescent antibodies to determine their cell type of expression or subcellular localization. Having the potential to overcome the limitations of both fluorescent secondary antibody and HRP labeling methods, the alkaline phosphatase (AP) immunolabeling method uses multiple alkaline phosphatase enzyme tethered to the secondary antibody to generate a fluorescent polymer deposit *in situ* for immunolabeling of specific protein antigen. The AP method has been used in histological analysis of non-brain tissue^*8*^. However, immunolabeling of proteins in the brain tissue present challenges due to the complexity in cell composition, high lipid content, and the presence of fine structures such as synapses where certain proteins are concentrated. The utility of AP immunolabeling method has not yet been demonstrated in the brain.

In this study, we provide the first demonstration of the capabilities of AP immunolabeling method to detect a wide range of antigens in the brain cells at the cellular and subcellular resolution.

Through side-by-side comparisons between standard Alexa-Flour antibody, HRP, and AP immunolabeling methods, we show that the AP method provides highly sensitive and specific labeling of the FLT3 kinase in the developing and adult mouse brain samples. The ability for AP labeling to be used in conjunction with regular immunofluorescence method enables co-staining with neuronal and glial subtype markers, which reveals a neuron-specific pattern of FLT3 expression in the brain. We then applied the AP method to achieve immunolabeling of Kir2.1, an inward-rectifying potassium channel that is weakly expressed in the brain, within the mouse, pig, and human brain tissue samples. We further investigated the extent to which AP immunolabeling can be used to detect neuronal antigens enriched in microscopic subcellular structures such as the synapse. Taken together, our results show that the AP polymer-based immunolabeling is a versatile methodology applicable for detecting various brain proteins in tissue or cell culture preparations from different animal species with high sensitivity and synaptic resolution.

## Results

### Establish the spatial-temporal pattern of FLT3 expression during brain development with AP immunolabeling method

We first seek to establish the expression pattern of FLT3 in brain tissue as a foundation for mechanistic investigation of FLT3 signaling in the brain. We focused on the cerebellum tissue since previously published results show enriched RNA expression in this brain region^*5*^. We adapted a AP-polymer histochemistry methodology that is used in histology analysis of peripheral tissue but not on the brain tissue^*8*^. Comparing to the standard fluorescent secondary antibody-based immunostaining method that failed to detect a consistent pattern of FLT3 expression, AP immunolabeling selectively labels FLT3 protein in the mouse brain slices in a pattern comparable with the HRP method which develops monochromatic deposit (**Fig. 1A-B**). Interestingly, in AP-labled cerebellum brain slices the granular layer and molecular layer can be readily distinguished by their different Flt3 labeling pattern, whereas such difference is mostly diminished in the HRP-labeled tissue, presumably due to the lower spatial resolution of the HRP labeling method. Quantitative comparison showing substantially improved FLT3 labeling sensitivity and signal-to-noise ratio (**Fig. 1C**).

**Figure 1.**
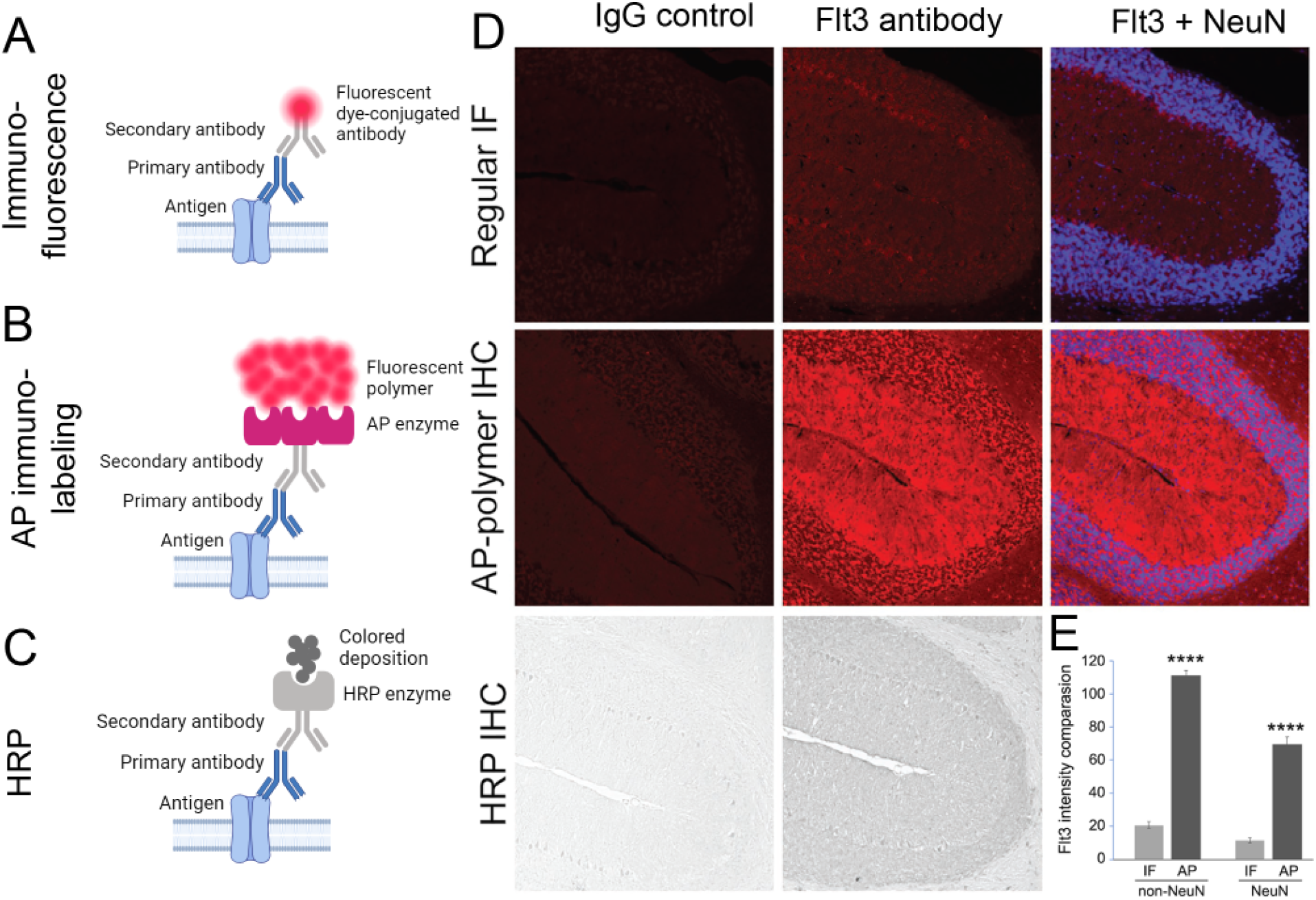
Visualizing Flt3 protein by different staining methods. Adult mouse cerebellum sagittal sections (cryostat, 10 µm in thickness) were immunostained by three different methods in order to visualize a weakly expressed protein Flt3. In the regular immunofluorescent staining (Regular IF in top panel, IF in graph). Flt3 was rarely seen in both molecular layer (ML) and granular layer (GL) with weak signals in the Purkinje cell layer (arrow heads in image, PCL in graph). However, in the alkaline phosphate-based histochemistry method (AP-polymer IHC in middle panel, AP in graph), Flt3 was seen by strong fluorescent positive signals, with the intensity > 5-fold higher than that from the regular IF in the ML+PCL area, almost 7-fold higher in GL area (graph). HRP-based histochemistry method (HRP IHC in the bottom 2 images) showed clear positive signals, but the staining was not able to show the intensity differences in ML and GL. Comparing two histochemistry methods, besides the differential signals in different areas, one prominent advance of AP-polymer IHC is that the staining is fluorescent, which provides the base for the hybrid method with other fluorescent staining. In all of the 3 staining methods, the staining control (Ctr) were processed exactly the same except applying the same amount of rabbit IgG replacing the Flt3 rabbit primary antibody (Flt3). Scale bar: 200 µm. ^***^P < 0.001 (AP vs. IF). One-way ANOVA followed by Bonferroni test. Mean ± SEM, N = 16 – 18 image areas from 3 individual staining with each method.

Importantly, the immunolabeling conferred by the AP method is fluorescent in the Cy3 channel but does not interfere with other fluorescent channels. Therefore, a hybrid protocol that use AP immunolabeling in conjunction with standard immunostaining methods enables multiplexed co-labeling of various antigens. Our results reveal a previously uncharacterized pattern of FLT3 protein expression in the mouse cerebellar tissue: The expression of FLT3 is exclusively in neurons, with no overlap with GFAP+ astrocytes or Iba1+ microglia (**Fig. 2A**). Interestingly, the expression of FLT3 seems to be especially high in the Purkinje cells that stain positive for PV and Calbindin, but not in the NeuN+ granular cells (**Fig. 2B**).

**Figure 2.**
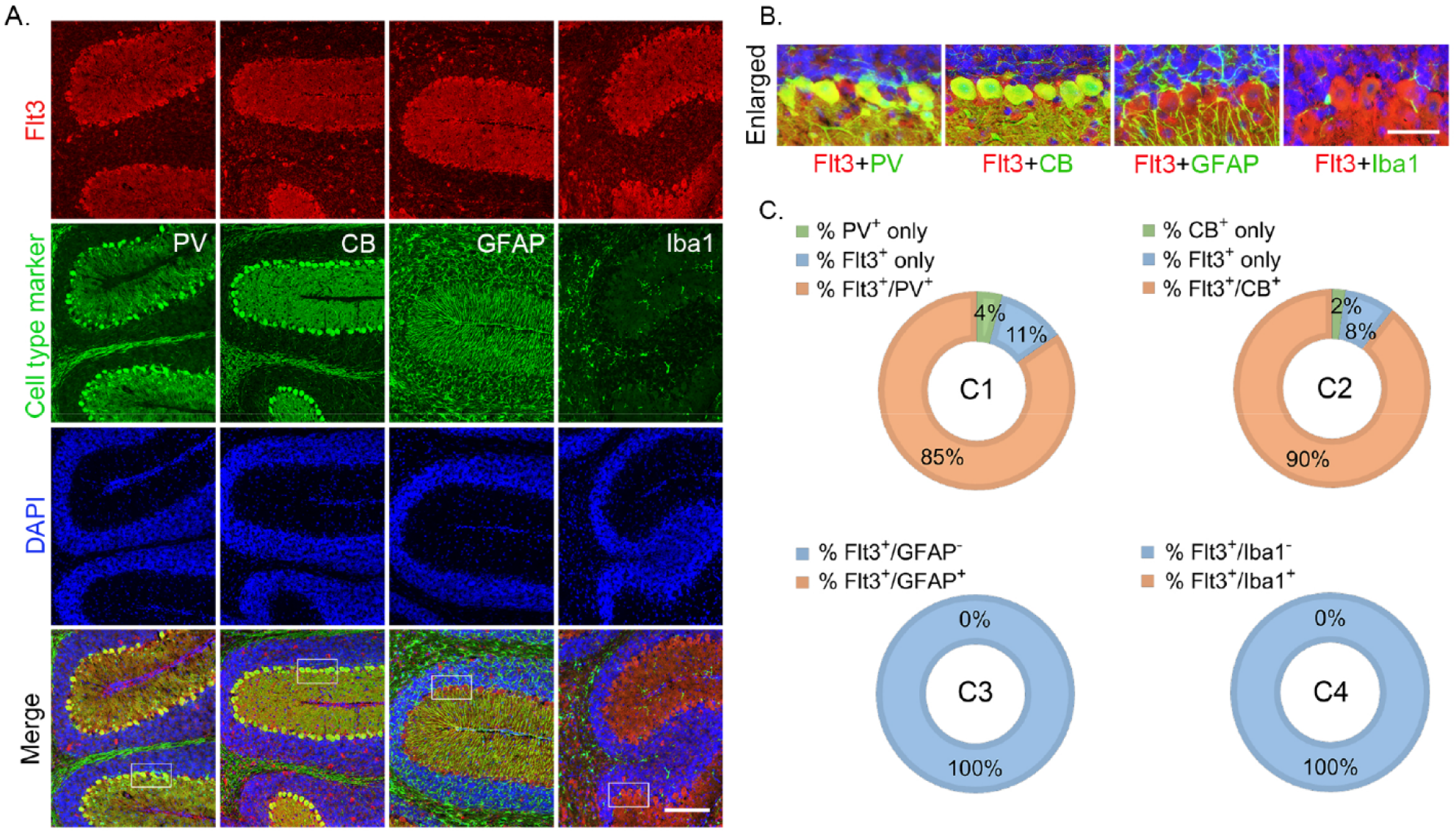
Flt3 is specifically expressed in inhibitory neurons in cerebellum. We created a hybrid method by combining AP-polymer IHC with other regular immunofluorescent staining to visualize the cell types and intracellular location of Flt3 protein. Postnatal D14 mouse brain sections were used for the staining. Flt3 (red) was visualized by AP-polymer IHC and other cell markers (green) were used as the regular IF staining. Flt3 was co-localized with the inhibitory neuronal markers parvalbumin (PV) and calbindin (CB) in the Purkinje cell layer, but not with the glial cell marker GFAP or the microglial cell marker (Iba1) (A and enlarged images B). In the PCL, 85% cells were Flt3 positive (Flt3+/PV+), 4% are PV positive but Flt3 negative (PV+ only), and 11% cells are Flt3 positive and PV negative (Flt3+ only, C1). About 90% cells are Flt3+ and CB+ (Flt3+/CB+, C2). None of Flt3+ cells were GFAP+ (C3) or Iba1+ (C4). Scale bars in A:200 µm, B: 50 µm. Mean ± SEM, N = 11 – 12 cerebellum images.

We further utilized the AP labeling method to investigate the temporal trajectory of FLT3 expression during mouse brain development, and found an overall increase in FLT3 immunoreactivity in cerebellum tissue samples collected at postnatal day 7 to two months of age, indicating an increases in FLT3 protein expression during cerebellar development (**Fig. 3A-B**). Importantly, the subcellular expression pattern also switches from predominantly cytosolic during early stages of brain development to displaying substantial dendritic portion in the mature brain (**Fig. 3C**). Taken together, AP immunolabeling of FLT3 not only establishes for the first time the brain region and cell type enrichment pattern that is in high agreement with single-cell sequencing data, but also enables the in situ detection and quantification of protein localization throughout developmental stages to reveal a cytosol-to-dendrite change in FLT3 subcellular localization in cerebellum Purkinje cells. These findings provide the foundation for further investigation to elucidate the roles of FLT3 in brain health and disease.

**Figure 3.**
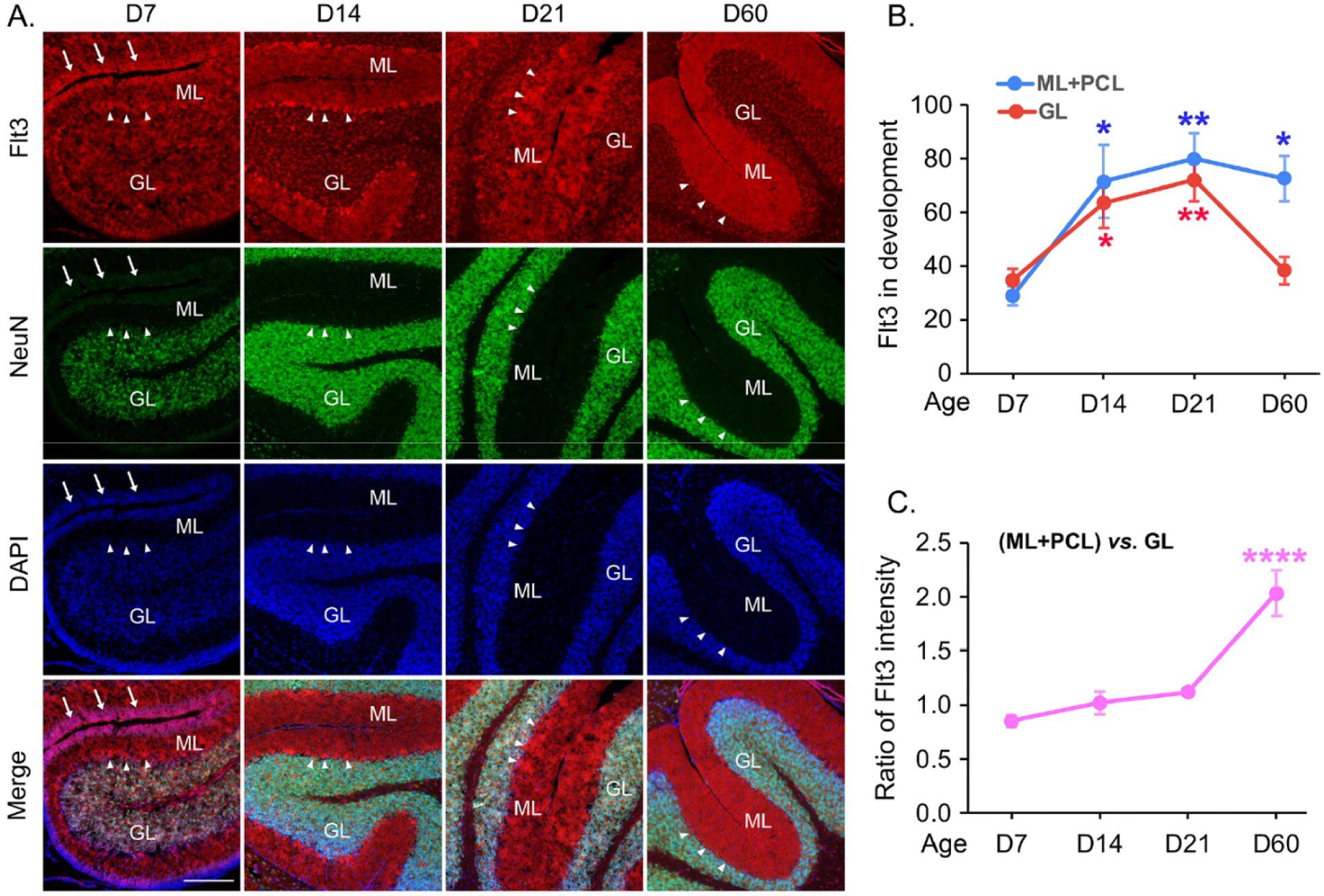
Flt3 increases in the cerebellum development and shifts from GL to ML. C57 mouse brains were harvested from postnatal D7, D14, D21, and D60, and their cryostat sections were co-stained for Flt3 and NeuN with the hybrid staining method (A). At D7, Flt3 was expressed at the low level both in ML, GL, PCL (arrow heads), and in the epineurium layer that was NeuN negative (arrows). At D14, Flt3 expression dramatically increased in both in GL, PCL and ML (A, B), and most of Purkinje cells were filled with Flt3 (A and in Figure 2A, B). At D21, the increased Flt3 started to shift from GL to ML (A), and at D60, most of Flt3 was stained in ML. The shift is substantial and dramatic, showing as the ratio of (ML+PCL) vs. GL (C). Scale bar in A: 220 µm. ^*^P < 0.05, ^**^P < 0.01 in comparison with their own D7 data in B. ^****^P < 0.0001 in comparison to D7 data in C. One-way ANOVA followed by Dunnett test. Mean ± SEM, N = 9 – 12 pairs of sections from 3 – 4 individual mouse brains at each time point.

### AP immunoabeling of potassium channel Kir2.1 in mouse, pig, and human brain tissue samples

We further investigated to which extent AP method can be applied to immunolabel membrane proteins, such as ion channels, that are expressed at low levels in the brain. Kir2.1 is a voltage-gated inward-rectifying potassium channel that plays a critical role in regulating the resting membrane potential in neurons. The expression level of Kir2.1 is low and under strict control to maintain neuronal excitability at the optimal level. Moreover, the cell-type specific pattern of Kir2.1 expression in the brain is not clear. We compared the immunostaining results of Kir2.1 in mouse brain tissue using standard or AP immunolabeling methods, and found a substantially improved labeling sensitivity and S/N ratio using the AP method (**Fig. 4A**).

**Figure 4.**
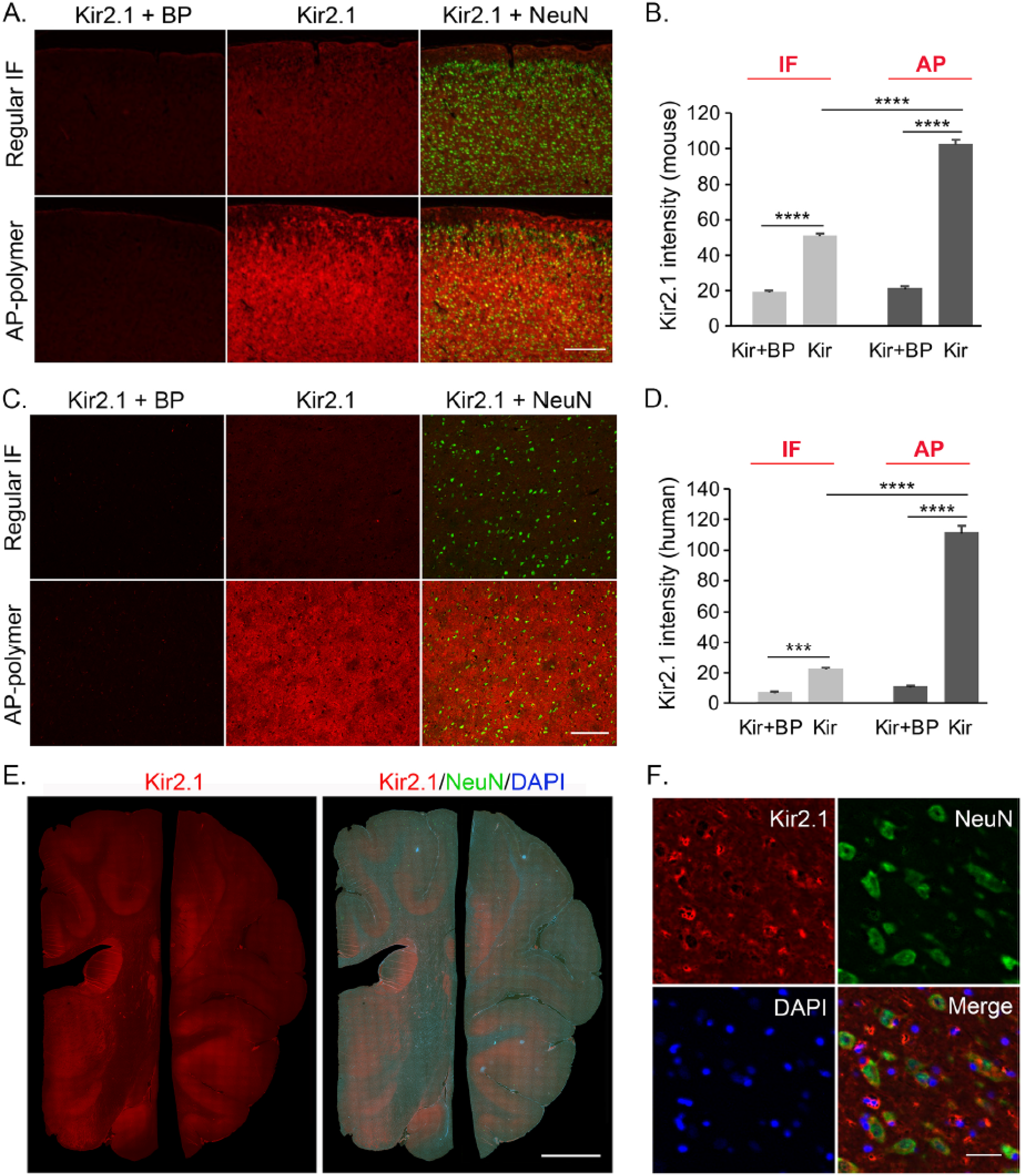
Hybrid staining method is suitable to use in different species brains. Kir2.1 is one of the potassium channels on the cell membrane and sometimes not easy to see clearly by the regular immunofluorescent staining. The regular immunofluorescent staining (regular IF in A, C; IF in B, D) and the hybrid staining (AP-polymer in A, C; AP in B, D) were compared respectively in mouse cortex (A, B) and human cortex (C, D, after years of formalin fixation). All staining was to co-label Kir2.1 and NeuN, and the staining control was processed exactly the same with others except the Kir2.1 primary antibody was incubated with its blocking peptide (Kir2.1+BP) 30 minutes before applying onto the sections. Quantitation of the staining intensity vs. its own control was summarized in B and D. The hybrid method was also used in pig brain (E, stitched image) and the enlarged cellular images showing in F. Scale bars in A: 450 µm; B: 400 µm; E: 2.5 mm; F: 30 µm. ^***^P < 0.001, ^****^P < 0.0001. Mean ± SEM, N = 15 mouse cortex areas were analyzed in B, N = 31 – 41 human cortex areas were analyzed in D. One-way ANOVA followed by Bonferroni test.

**Figure 5.**
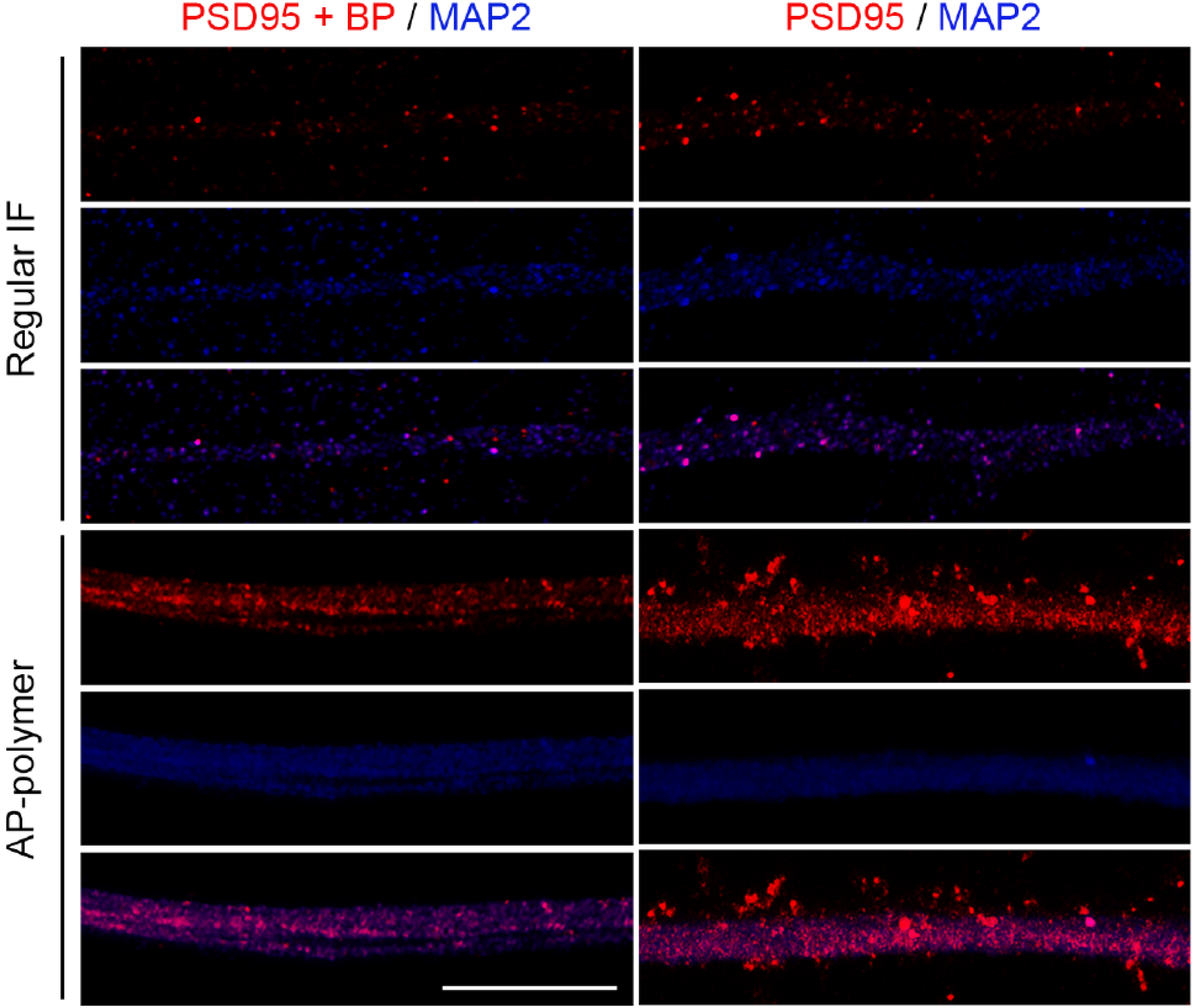
Hybrid staining method is capable for fine structures in dendritic spines. Human iPSC-neurons were cultured for 5 months and stained with PSD95 (red) and MAP2 (blue) antibodies by the hybrid staining method (AP-polymer). The regular immunofluorescent staining was used for a comparison (Regular IF). The staining control was using PSD95 antibody pre-incubated with its blocking peptide (PSD95+BP), to replace the PSD95 antibody. Images were taken from Zeiss LSM 980 airyscan 2 confocal microscope (63x object). While very few signals from the regular IF staining, the hybrid method provides much more and brighter dendritic spine-like button staining. Scale bar: 10 µm.

We further performed analysis of mouse brain slices in which Kir2.1 protein are AP immunolabeled, while various brain cell types including neurons, astrocytes, and microglia are labeled by NeuN, GFAP, and Iba1 antibodies, respectively. We found that the majority of Kir2.1 immunoreactivitiy surrounding NeuN+ neuronal nuclei (**Fig. 4B**). We then validated our finding of neuron-enriched Kir2.1 expression pattern in the mouse brain slices to pig and human cortical tissue sections using the same immunolabeling procedures (**Fig. 4C-D**). Similar labeling of Kir2.1 can be achieved.

### AP immunoabeling of PSD95 in the synapses within human stem cell-derived neurons

Having established that the AP immunolabeling method can be used to label protein localization at cellular resolution, we further analyzed to what extent the fine subcellular structures such as the synapses can be precisely labeled with the AP method. We employed cultured human stem cell-derived neurons that are widely-applied in disease model and in drug screening^*7, 9*^. Our previous work report the functional maturation of synapses in human stem cell-derived neurons with the support of astrocytes^*10*^. We further looked at synapses in human iPSC-derived neurons. Utilizing MAP2 antibody to label the neuronal dendrites and PSD95 antibody with either regular antibody or AP immunolabeling method to mark the postsynaptic density. Comparing to standard immunostaining method, AP method can clearly label a critical component of the postsynaptic density PSD95 with high sensitivity. Taken together, our results indicate that the AP immunolabeling method can be used to investigate fine structures such as the synapses, therefore presents a useful tool for neuroscience research for studying the cellular and subcellular distribution of neuronal antigens.

## Discussion

In this study, we demonstrated the feasibility and established an optimized workflow of applying a highly sensitive AP-based immunolabeling method in brain tissue to label a variety of proteins that play important roles in neuronal function. We demonstrated successful immunolabeling of the developmental trajectories in protein localization, at cellular and subcellular resolutions, of receptor tyrosine kinase FLT3, potassium channel Kir2.1, and post synaptic density scaffolding protein PSD95. The versatile AP immunolabeling method can work with a wide range of samples including mouse, pig, and human brain tissue samples, as well as human stem cell-derived neurons. Our results uncover that FLT3 expression in the cerebellum tissue is highly enriched in neurons, especially in GABAergic Purkinje neurons, and that during cerebellum development the localization of FLT3 shifts from being primarily cytosolic to enriched in the dendrites. These results complement our group’s previously reported findings showing that FLT3 signaling modulates neuronal gene expression and function, and provides a foundation for further studies to elucidate FLT3 signaling cascade in the brain.

To further illustrate the wide applicability of AP method for immunolabeling low-abundance antigens in brain tissue, we used AP-immunolabeling method to establish a neuron-enriched expression pattern of Kir2.1 in the mouse brain, a feature that is evolutionarily conserved in the brain of pigs and human. This result form a cellular basis for the development and application of therapeutics targeting Kir2.1. An important and thriving field of neuroscience research is focused on synapses, subcellular structures that contain a multitude of proteins and are at the limit of light microscopy detection. Our work show that the AP immunolabeling can be used in cultured human stem cell-derived neurons to label the PSD95 protein in synaptic structure. The capability to resolve synapse-scale structures open many opportunities for applying the AP immunolabeling method to

In summary, the versatility of the AP immunolabeling methods, and its compatibility with standard immunohistology and immunocytochemistry methodologies, substantially expands the use case of this method for investigating proteins in the brain. Our results highlight the power of applying in situ protein detection method such as AP immunolabeling to validate and functionalize RNA sequencing results and yield new insights into the protein products of the genes of interest.

## Methods

### Primary and stem cell-derived neuron culture

Primary mouse cortical neurons were obtained through dissection of neonatal mouse brains and enzymatic dissociation of the cortical tissue. The cells were seeded at a density of roughly 100,000 cells per well into 24-well plates containing poly D-lysine coated glass coverslips.

Human embryonic stem cells (hESC) derived neural progenitor cells (NPC, generated from WIBR1 hESC line) were seeded at 10,000 cells per well onto glass coverslips pre-seeded with an astrocyte feeder layer^*10*^.

### Immunohistochemistry tissue sample preparation

Mice were euthanized and perfused with 4% PFA at all required ages and their brains were post-fixed in PFA for 2 days, then changed to 30% sucrose until cryostat sectioning. The O.C.T. embedded brain samples were cut at 10 µm in thickness with Leica cryostat machine and directly mounted the sections onto the VWR superfrost glass slides. Paraffin-embeded pig coronal sections were acquired from Dr. Jianhua Qiu in Mannix lab. Formalin-fixed human brain tissues were acquired from the BCH pathology department and processed in the BI Pathology Core for paraffin-embedding and sectioning (5 µm in thickness). Two brain sections were mounted on each slide.

### Immunolabeling of tissue sample

Three different immunostaining methods were used in this work: the regular immunofluorescent staining, HRP-based histochemistry, and the AP-based histochemistry hybrid with the regular fluorescent staining.

1. Regular immunofluorescent staining The cryostat brain sections were heated on a heating plate for 20 min (37 - 42°C) to dry out the wetness from the freezer, and washed off O.C.T. in Tris buffered saline (TBS). The pig and human brain paraffin-embedded sections went through xylene and a series of different concentrations of ethanol and washed in TBS. All brain slides went through antigen retrieval at 95-100°C for 10 minutes (Vector Laboratories, H-3300- 250), then incubated with the blocking buffer (5% serum + 2% BSA in TBS-T (0.1% Tween-20)), primary and fluorescent secondary antibodies, then coverslipped with DAPI-fluoromount mounting medium (SouthernBiotech, 0100-20).
2. HRP-based histochemistry Similar to the regular staining, the brain section slides went through antigen retrieval and incubated with 0.1% H2O2 to inactivate the endogenous HRP. After blocking, one section on each slide was incubated with Flt3 primary antibody at 4°C overnight (Abclonal, A12462, 1:200) and the other section on the same slide was used as the staining control which used the same amount of rabbit IgG to replace the Flt3 rabbit antibody. On the second day, after the TBS-T wash, the slides were incubated with goat anti-rabbit biotinylated secondary antibody (Vector Laboratories, BA-1000, 1:500) followed by VECTASTAIN ABC Reagent (Vector Laboratories, PK-6100) and developed in freshly made DAB peroxidase substrate solution (Vector Laboratories, SK-4100). Monitor the development under the microscope until satisfied by comparing with the control section on the same slide. The stained slides went through a series of ETOH and Xylene, coverslipped with Cytoseal.
3. AP-polymer histochemistry hybrid with regular fluorescent immunostaining For the co-staining of Flt3 (rabbit) with NeuN (mouse), GFAP (chicken), Iba1 (goat), parvalbumin (chicken), calbindin (chicken) in brain sections, we use AP-polymer-based histochemistry to boost Flt3’s signal and co-staining with other cell specific markers. The brain slides were processed the same as in the above HRP-IHC method except using H2O2, until adding the secondary antibody. Instead of biotinylated secondary antibody in HRP histochemistry, the horse anti-rabbit AP-conjugated secondary antibody (Vector Laboratories, Cat#: MP-5401) was used for 10 minutes at room temperature. After the brief wash with TBS-T and TBS, develop AP with the substrate solution (Vector Laboratories, Cat#: SK-5105), monitor the color under the microscope, and stop the reaction by immediate TBS wash. AP-polymer histochemistry produced the red fluorescent precipitance (wavelength similar as Alexa 594). The brain slides were continued to be incubated with other fluorescent secondary antibodies for the co-stained fluorescence (e.g. Alexa 488, 647, etc.). There are several critical points to make a successful co-staining with this hybrid method: (1) After the AP development, no detergent buffer is allowed to use in the following co-staining procedures, since the detergent destroys the precipitate pattern from the AP histochemistry. (2) Always have a negative control with the same amount of IgG as the to-test primary antibody on the same slide. (3) Similar to the HRP histochemistry, endogenous AP should be considered. Heating slides (like antigen retrieval) is a good way to inactivate the endogenous AP. If the staining protocol does not include heating, AP inhibitor (Levamisole) may be considered.

### Imaging and quantification

With our strict principle for the control setting in each staining of every single slide (two sections on one slide; using same amount of IgG to replace the antibody; using blocking peptide to pre-incubate with antibody, exact same development time, etc.), images from the positive staining and the control staining were always taken with the exact same setting. In many circumstances, the single channel images and other co-stained channel images were first merged together with Image J in order to draw the exact areas for the quantification, then split the colors to only choose the target channel for the intensity measurement. On each slide, the measured intensity data were first normalized by its own staining control data from the other section on the same slide, then being included in the groups. The detailed sample numbers depended on the different experiments and described in each figure legend.

### Statistics

Described in each figure legend.

## Author contributions

Y.Y and X.T., designed research; Y.Y., K. H., J. K., I. H. performed research; Y. Y., K. H., and I. H. analyzed data; Y. Y. and X.T. wrote the paper.

## Funding

This work was supported by the SFARI Bridge to Independence grant, Charles Hood Foundation, SYNGAP Research Fund, PTEN Research fund, RSZ TNC fund, and NIH R01NS138758 awarded to X.T. We would like to thank the IDDRC Cellular Imaging Core for their support, funded by NIH P50 HD105351 and supported by NIH grant #S10OD030322.

## Acknowledgement

We graciously thank Dr. Jianhua Qiu and Dr. Rebeckah Mannix from Boston Children’s Hospital for sharing pig brain tissue, and thank the BCH pathology department for sharing the human postmortem brain samples.

## Notes

### Competing Interest Statement

The authors have declared no competing interest.

